# Sero-epidemiology of Rift Valley fever virus in ruminant livestock in The Gambia

**DOI:** 10.1101/2025.03.14.640388

**Authors:** Essa Jarra, Yahya Beyai, Alphonse Mendy, Ousman Ceesay, Abdou Ceesay, Kebba Daffeh, Daniel T. Haydon, Sarah Cleaveland

## Abstract

Rift Valley fever (RVF) virus is a globally significant zoonotic pathogen primarily transmitted by mosquitoes, causing abortion storms and neonatal mortality in ruminant livestock and haemorrhagic encephalitis in humans. Although localized RVF outbreaks occur frequently in the West Africa Sahel, the complex interplay between vectors, ruminant hosts, human behaviours, and ecoclimatic factors driving these outbreaks remains unclear. In 2022, we conducted a cross-sectional study across The Gambia that collected serum samples linked to questionnaire data from 202 randomly selected livestock-owning households to estimate population-level seroprevalence, investigate the patterns of, and identify risk factors associated with RVFV seropositivity in ruminant livestock. Design adjusted seroprevalence estimates were 36.8% (95% Confidence Interval [CI]: 31.6–40.0) for cattle (n = 1416), 2.1% (95% CI: 0.5–3.7) for goats (n = 1101) and 4.3% (95% CI: 1.2–7.4) for sheep (n = 1085) using the IDScreen® RVF c-ELISA assay. The intracluster correlation coefficients for RVFV seropositivity at the herd level were 0.13 (95% CI: 0.09–0.18) in cattle and 0.22 (95% CI: 0.06–0.31) in small ruminants on the logit scale. Risk factors associated with RVFV seropositivity were investigated using a LASSO regression, with transhumant movement to the Gambia river valley, increasing age, introduction of new ruminants, and proximity to water identified as most strongly associated with seropositivity. By fitting the serologic data to a catalytic force of infection (FOI) model, age-independent FOI estimates of 0.12 (95% Credible Interval [CrI]: 0.10–0.16) for cattle and 0.013 (95% CrI: 0.01–0.02) for small ruminants were obtained. The observed patterns of RVFV seropositivity indicate widespread and endemic viral circulation, with livestock management practices and the Gambia river floodplains likely sustaining transmission. The high seroprevalence in cattle raises concerns that sporadic outbreaks may have occurred unnoticed or unreported, underscoring the critical need for comprehensive One Health surveillance to identify potential clinical cases.

## Introduction

Rift Valley fever (RVF) is a vector-borne zoonotic disease that poses significant threats to public and animal health, as well as the socioeconomic stability of livestock-dependent communities [1]. The disease is caused by the Rift Valley fever virus (RVFV), a single-stranded RNA virus of the genus *Phlebovirus* [Family *Phenuiviridae*] [2]. Since its identification in Kenya in 1931, RVFV has triggered major outbreaks across sub-Saharan Africa, and increasingly beyond, with outbreaks recorded in the Arabian Peninsula and the Comoros Archipelago [3]. Its expanding epidemiological footprint is attributed to factors such as both legal and illegal livestock trade, and the encroachment of RVFV-competent vectors into new habitats [4, 5].

RVF outbreaks in ruminant livestock are often devastating, characterized by “abortion storms” affecting 40–100% of pregnant ruminants, and substantial neonatal mortality [5]. In humans, the disease typically manifests as a mild, flu-like illness, but 1–3% of cases develop severe haemorrhagic syndrome with multi-organ dysfunction and fatality rates up to 50% [4, 6]. Over 30 mosquito species, including those from the *Aedes, Culex*, and *Mansonia* genera are capable of transmitting RVFV, while other dipteran species may contribute to mechanical transmission [7, 8], reflecting its ecological adaptability. Infections in animals primarily occurs through mosquito bites, though direct exposure to infectious abortion tissues and fluids can also facilitate transmission [5]. Humans are predominantly infected through contact with viraemic livestock, contaminated products (e.g. raw milk and meat), or aerosols during outbreaks [9-11]. Notably, human-to-human transmission has not been documented, except for rare cases of *in utero* transmission [12].

RVF epidemiology is characterized by two distinct but overlapping cycles: large outbreaks interspersed with prolonged inter-epidemic periods (IEPs) marked by low-level, cryptic virus circulation [13, 14]. In East Africa, large outbreaks are strongly linked to climatic and ecological drivers, particularly heavy rainfall associated with the El Niño-Southern Oscillation (ENSO) phenomenon [15], which promotes mosquito population surges. These outbreaks recur at an average IEP of 7.9 years (range: 3–17 years) in Tanzania [16] and 9 years (range: 2–21 years) in Kenya [17]. In contrast, large outbreaks in West Africa are geographically confined to Mauritania and Senegal, occurring every 2–5 years and sometimes unrelated to heavy rainfall [1, 18, 19]. The most recent RVF outbreak in humans and animals was in 2022 in Mauritania, with 47 confirmed human cases and 23 deaths from haemorrhagic syndrome [20].

In the RVF-endemic Ferlo region of Senegal, rain-fed temporary ponds serve as critical breeding groups for *Aedes vexans* and *Culex* spp. and attract large numbers of ruminants, creating favourable conditions for RVFV transmission [21, 22]. Risk factor analyses of RVFV seropositivity in animals during the IEP have identified various livestock management risk practices [23-25]. A particularly important factor in West Africa is the role transhumant and nomadic movement of ruminant livestock to reach watering and grazing points [22], play in creating a “source-sink” dynamic wherein RVFV can persist and spread across large areas.

Despite established links between climatic cycles, vector ecology and RVFV transmission, significant knowledge gaps remain regarding the mechanisms supporting viral maintenance during the IEP [26-28]. While transovarial transmission of RVFV in *Aedes mcintoshi* has been demonstrated in East Africa [29], robust empirical evidence supporting this mechanism in West Africa is lacking, particularly for *Aedes vexans*, the dominant *Aedes* spp. implicated in RVFV transmission in this region. Alternative hypotheses suggest RVFV survival in overwintering *Culex* spp. mosquitoes [30, 31] or limited roles of wildlife reservoirs [18, 32], as mechanisms sustaining viral circulation between large outbreaks. Regardless of the exact mechanism, there is sustained low-level RVFV circulation during the IEP, with increasing reports of livestock seroconversion outside major outbreaks across sub-Saharan Africa [33-35]. The incomplete understanding of RVF dynamics highlights the need for further research on the contributions of different potential risk factors in the dry Sahelian zone.

RVF was first reported in humans and small ruminants in The Gambia in 2002 [36]. The absence of subsequent outbreak reports raises questions about whether RVFV continues to circulate undetected in the livestock populations. However, a fatal human RVF case of a resident of The Gambia was reported in 2018 [37]. Indeed, limited surveillance capacity in remote livestock production areas where the risks of infection are likely to be highest [38], may miss low-level, endemic transmission that could be occurring within the livestock populations. The Gambia presents a unique setting for understanding RVF epidemiology in West Africa due to its proximity to well-known endemic countries of Mauritania and Senegal, its large ruminant livestock populations, and distinct geography spanning along the Gambia river. However, very little is known of RVFV epidemiology in The Gambia and the factors associated with exposure to infection.

This study aimed to estimate population-based RVFV seroprevalence and force of infection, identify livestock management practices associated with RVFV seropositivity among ruminant livestock, and examine spatial variations in seroprevalence across The Gambia.

## Materials and Methods

### Study area

The study was conducted across the six administrative regions of The Gambia: Central River Region-North (CRR-N), Central River Region-South (CRR-S), Lower River Region (LRR), North Bank Region (NBR), Upper River Region (URR) and West Coast Region (WCR) (Figure 1). The country is a narrow enclave of Senegal and occupies a transitional ecoclimatic zone between the Sahara Desert to the north and the Guinean savanna to the south [39, 40]. The primary geographical feature of the country, the Gambia river, runs for 450km through its length, providing essential freshwater resources for agriculture and supporting a range of ecosystems, including mangrove swamps and floodplains that are critical for livestock grazing.

**Figure 1:**
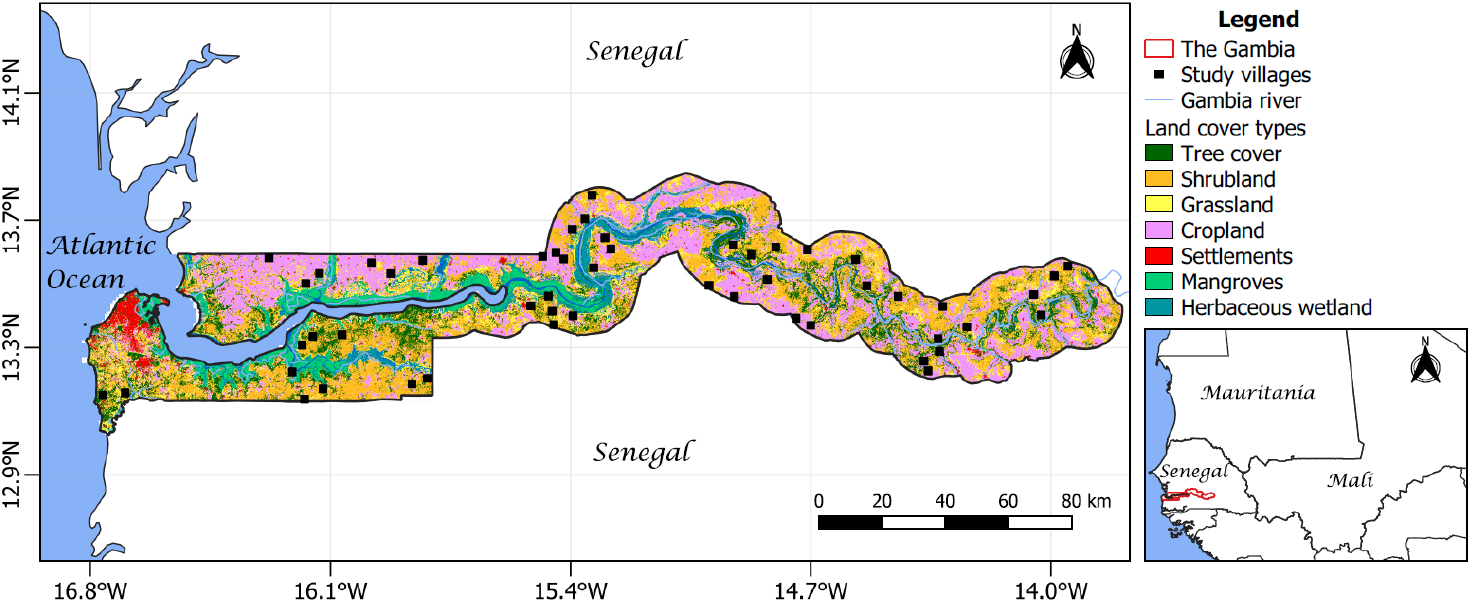
Map of The Gambia illustrating the location of villages selected for household surveys and ruminant sampling. The Gambia river runs along the length of the country. Land cover data: European Space Agency (ESA) WorldCover project 2021 / Contains modified Copernicus Sentinel data (2021) processed by ESA WorldCover consortium. Map was created in QGIS version 3.28.3, and the publicly available shapefiles were downloaded from the Database of Global Administrative Boundaries - GADM [www.gadm.org].

The climate is characterized by a unimodal wet season from July to October, with annual precipitation ranging from 650 to 1200 mm [41]. Rainfall during this period can be intense with high humidity, whereas the dry season (November–June) is marked by arid conditions with temperatures reaching 40°C. The topography is predominantly low-lying and flat, comprising a mixture of grasslands and freshwater floodplains, interspersed with low hills and shallow depressions that experience seasonal flooding. During the wet season, most of the grasslands are transitioned into cultivated land for rain-fed agriculture. Wildlife in the region includes various species of monkeys, baboons, and antelopes. Data collection for this study coincided with the peak dry season (February–May).

Livestock production is crucial to the Gambian economy and for creating livelihood opportunities, contributing approximately 30% of agricultural GDP and 10% of national GDP [42]. The 2016 National Livestock Census [43] reported populations of 292,837 cattle, 172,662 sheep and 328,336 goats. The predominant livestock management system is traditional, extensive, and integrated with crop agriculture. Cattle are herded daily for 6–8 hours to graze on unimproved communal grasslands, and are tethered individually on the outskirts of villages at night [44]. In contrast, small ruminants (sheep and goats) are usually managed separately from the cattle and confined within residential compounds in enclosures made of mud bricks or wooden planks, with roofs of thatch or corrugated sheets.

### Ethical clearance

The study protocol received ethical approval from the National Agricultural Research Institute of The Gambia [Approval Reference: NARI/DLS/01/(01)] and the University of Glasgow School of Veterinary Medicine Research Ethics Committee (Reference: EA47/21). Approval for interactions with human participant were obtained from The Gambia Government/MRCG@LSHTM Joint Ethics Committee (Project ID/Review Ref: 26750) and the University of Glasgow College of Medical, Veterinary and Life Sciences Ethics Committee (Project No: 200210057).

Community engagement meetings were held to introduce the study, explain sampling procedures, and obtain informed consent. Participants provided written consent or a thumbprint as a substitute in cases of illiteracy. Consent procedures emphasized voluntary participation, anonymity, and participants’ right to withdraw. Collaboration with the local livestock assistants of the Department of Livestock Services (DLS), who are familiar with and trusted by the communities, facilitated recruitment of participants. All DLS livestock assistants received biosafety and ethics training prior to data collection. No human samples were collected for this study.

### Design and sampling strategy

The sampling frame for this cross-sectional survey included all cattle, sheep, and goats from 5,188 cattle-owning households recorded in the 2016 National Livestock Census [43]. The household list was stratified by administrative area. We fitted a Poisson distribution to the number of cattle-owning households in each region using the *fitdistr* function from the *MASS* package [45]. This allowed us to estimate the Poisson lambda parameter, representing the mean number of households to recruit in each region. Villages within each district (mean = 7, range = 5–9 per region) were selected using a two-stage cluster random sampling approach. A total of 52 villages were visited between February and May 2022. Up to four livestock-owning households per village were then randomly selected using the Livestock Census records. Data on livestock demographics, household characteristics, and management practices in the six months preceding the survey were collected through a pre-tested structured questionnaire on the Open Data Kit (ODK) platform [https://opendatakit.org] [46] on electronic tablets.

### Sample size and power analysis

A pilot analysis of biobanked cattle sera collected from 2019–2020 in The Gambia indicated RVFV seropositivity of 30% (n = 552, 95% Confidence Interval [CI]: 26.7–34.4). We conducted simulation-based power analysis using generalized linear mixed models (GLMMs) [47], to explore the impact of varying study design choices (number of ruminants, households, and villages) on statistical power based on the pilot data. This analysis indicated that sampling eight animals of each species (cattle, sheep, and goats) from 172 households would allow detection of 30% seropositivity with >90% power and ±7% margin of error for the household herd movement variable. A combination of haphazard and convenience sampling was used to select animals in each recruited household, and blood samples were collected via jugular venepuncture. Samples were allowed to clot at ambient temperature, centrifuged, and serum stored at -20°C until serological analysis.

None of the recruited households kept detailed records of the ages of their animals. Consequently, age was estimated based on the dentition of individual animals sampled [48, 49], incorporating information provided by owners and expertise of the livestock assistants for older animals with a full mouth of permanent incisors. Animals under six months old were excluded to prevent maternal antibody interference. Age was categorized into 1-year intervals with the midpoint of each age group used for analysis.

### Serological detection of RVFV antibodies in livestock

Serum samples were screened for anti-RVFV IgG antibodies using a competitive multispecies nucleoprotein ELISA (c-ELISA) assay (IDScreen® Rift Valley Fever Competition Multi-species, ID-Vet), according to the manufacturer’s instructions [50]. This assay was validated in 2021 at a regional veterinary reference lab in Senegal (LNERV), with 45 RVFV-positive and 133 RVFV-negative reference samples tested against the serum neutralization test which is considered the gold standard for RVFV antibody detection [51]. The sensitivity and specificity of the ID-Vet assay were estimated at 100% and 98.49%, respectively [52]. Serum samples with competition percentage (S/N % = (OD_sample_/OD_negative control_) × 100) ≤40% were interpreted as seropositive, those between 40%–50% were considered doubtful and those >50% were considered negative for RVFV IgG antibodies. For data analyses, all samples with a doubtful competition percentage were classified as seronegative. To test the robustness of our conclusions to changes to the classification threshold for seropositivity, the data analyses were repeated with doubtful c-ELISA results classified as seropositive.

### Data analyses

Household-level data on livestock management practices, abortion and mortality history, and disease control strategies were collected through a structured questionnaire administered to household heads or livestock owners. The survey data were cleaned in R version 4.2.2 [53], and individual animal serological results were merged. The final data set was stratified by species groups: cattle and small ruminants (goats and sheep). Means, standard deviations, and percentages were calculated using the *survey* package [54]. Data visualization was conducted with packages *ggplot2* [55] and *ggpubr* [56].

RVFV seroprevalence and 95% confidence intervals were calculated using the *svyciprop* function (method = “likelihood”) in the *survey* package [54], adjusting for unequal sampling using animal and herd-level weightings [25] – a herd being the total number of individuals of a species group (cattle and small ruminants) within a household. The animal-level weighting (ω_aj_) was calculated as the proportion of animals sampled from a given herd, while the herd-level weighting (ω_hk_) was calculated as the proportion of herds sampled within each village [25]. The overall weighting was the product of ω_aj_ and ω_hk_. Population-level seroprevalence estimates were also adjusted for the assay sensitivity and specificity based on validation data from LNERV, Senegal. The herd-level intracluster correlation coefficient (ICC) was individually calculated on the logit scale for each species group using the random effect variance of a null mixed model [57], to estimate the extent to which variability in seropositivity is due to between herd differences.

Univariable mixed-effects binomial models were initially fitted using the *glmer* function from the *lme4* package [58] in R, to screen associations between RVFV seropositivity and potential risk predictors/consequences of RVFV exposure. Random effects for district and village were included. To address multicollinearity and handle the large number of predictor variables collected during the survey (eighteen in total), variable selection was performed using a Least Absolute Shrinkage and Selection Operator (LASSO) regression [59]. We applied the glmmLasso algorithm (version 1.6.2) [60], which is based on a Lasso-type regularization in GLMMs to select key variables by shrinking some coefficients to zero, retaining those most strongly associated (non-zero coefficients) with RVFV seropositivity. A flowchart of the glmmLasso procedure is available in Supplementary Figure S1. Following Zhao et al. [61], a final logistic GLMM that includes only the selected (non-zero) variables was fitted to obtain unbiased coefficient estimates.

### Estimating species-specific RVFV force of infection (FOI)

FOI represents the rate at which susceptible animals in a population become infected, expressed as the probability of acquiring infection per unit time. In serological studies, it often reflects the transition from seronegative to seropositive status and is influenced by age, contact patterns, and exposure intensity to the pathogen [62]. We estimated FOI among cattle and small ruminants using different parametric models described in Hens et al. [62]: comparing a constant FOI model (Muench’s catalytic model [63]) to two age-dependent models: Griffiths’ linear model [64] and the Grenfell-Anderson’s quadratic model [65]. We assessed the suitability of each model for our serologic data across 1-year age groups using the Akaike Information Criterion (AIC) to compare model fits.

## Results

### RVFV seroprevalence and clustering

The study completed cross-sectional surveys in 202 livestock-owning households across 52 villages, sampling 3602 ruminant livestock (1416 cattle, 1101 goat, and 1085 sheep). Seroprevalence results for cattle and small ruminants are presented in Tables 2 and Table 3, respectively. Among the 202 households, 168 owned cattle and 165 owned small ruminants. Of these, 90% of cattle-owning households owned at least one seropositive cattle, whereas 31% of small ruminant-owning households had at least one seropositive small ruminant. The estimated ICC for RVFV seroprevalence between household herds were 0.13 (95% CI: 0.09– 0.18) in cattle and 0.22 (95% CI: 0.06–0.31) in small ruminants, calculated on the logit scale (see Supplementary Text).

After adjusting for the survey design and diagnostic assay, the overall RVFV seroprevalence in this study was 13.1% (95% CI: 11.0–15.3). The seroprevalence was 36.8% (95% CI: 31.6– 40.0) for cattle, 2.1% (95% CI: 0.5–3.7) for goats and 4.3% (95% CI: 1.2–7.4) for sheep. Seropositivity was significantly higher in cattle than small ruminants, 𝒳^2^ (df = 1, n = 3602) = 499.15, p < 0.001. The odds of RVFV seropositivity in cattle (reference species) was more than 90% higher than in goats (OR = 0.05, 95% CI = 0.03–0.07) and sheep (OR = 0.06, 95% CI = 0.04–0.09) in The Gambia. Although reclassifying all doubtful c-ELISA results (40% < S/N% ≤ 50%) as seropositive increased the adjusted RVFV seroprevalence to 45.3% (95% CI: 40.9– 49.7) for cattle, 5.9% (95% CI: 3.7–8.0) for goats, and 4.9% (95% CI: 2.1–7.8) for sheep, the core conclusions of our study remained unchanged.

The adjusted RVFV seroprevalence in cattle varied considerably across districts, ranging from 9.8% (95% CI: 3.7–16.0) to 57.5% (95% CI: 37.9–77.2), and from 0 to 69.8% (95% CI: 47.1–92.4) across villages. In contrast, the spatial variation of seroprevalence for small ruminants was smaller, ranging from 0 to 8.9% (95% CI: 6.3–11.55) across districts and from 0 to 29.0% (95% CI: 26.0–32.0) across villages. These variations in RVFV seroprevalence across study villages are illustrated in Figure 2.

**Figure 2:**
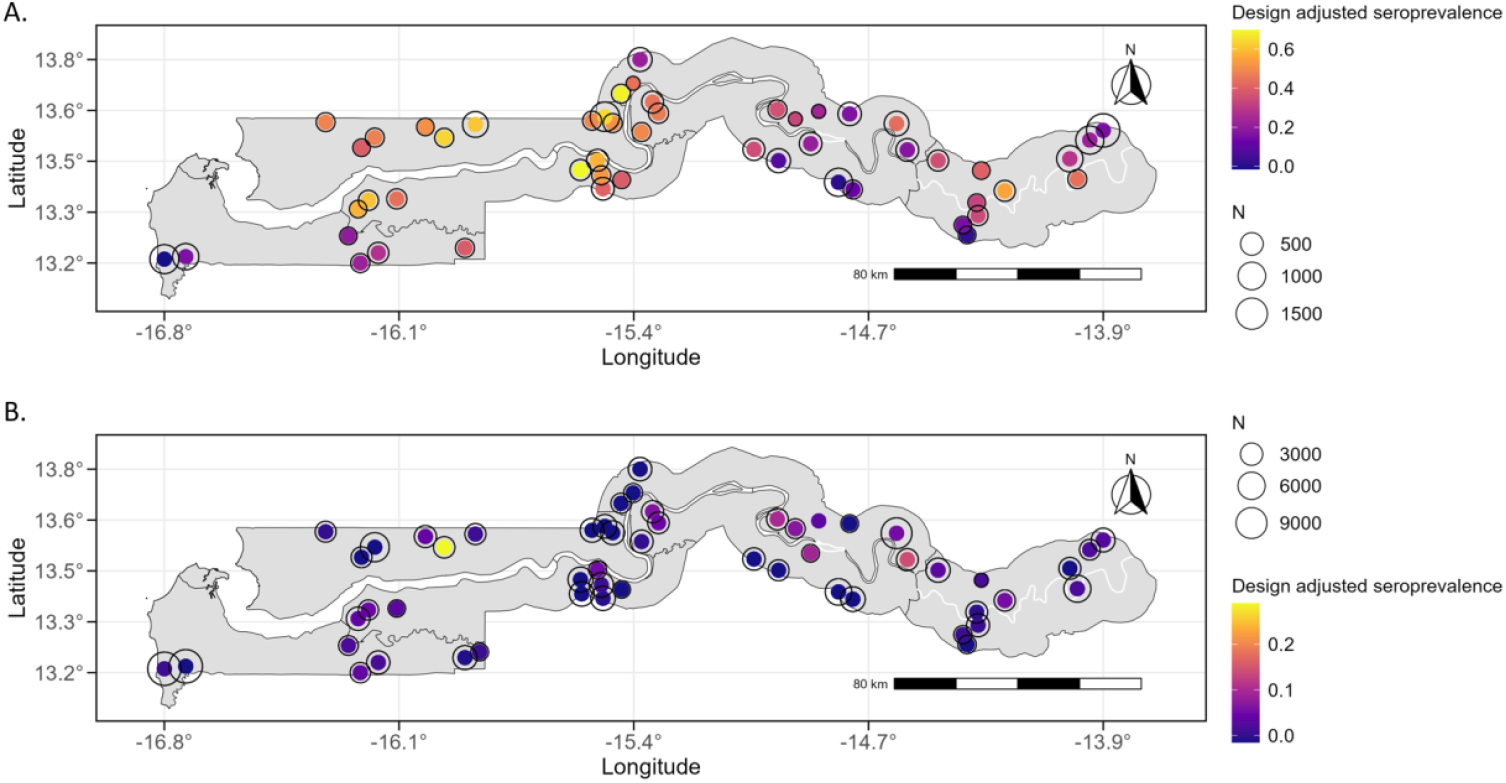
Adjusted RVFV seroprevalence in cattle **(A)** and small ruminants **(B)** overlaid with village herd population size (N) of each species group according to the 2016 Livestock Census.

### RVFV force of infection

The FOI for cattle and small ruminants in The Gambia was modelled using three different approaches. The AIC values indicated that the Grenfell-Anderson quadratic model [65] provided the best fit for our age-stratified serologic data (Table 1).

**Table 1:** Comparison of FOI model fits for the cattle and small ruminant populations using Akaike information criteria value (AIC). The lowest AIC is highlighted in **bold** for each species group.

| FOI Model | AIC |  |
| --- | --- | --- |
|  | Cattle | Small ruminants |
| Muench's model (constant FOI) | 104.390 | 36.999 |
| Griffiths' model (Linear FOI) | 62.117 | 33.908 |
| Grenfell-Anderson's model (Quadratic FOI) | **57.100** | **31.228** |

The cumulative FOI, derived by integrating age-specific FOI estimates from the quadratic model, was used to model RVFV seroprevalence. Predicted seroprevalence closely matched observed values across age groups of both cattle and small ruminants (Figure 3), with narrow confidence intervals indicating robust model performance. Age-independent annual FOI estimates using the Muench catalytic model [63] were 0.12 (95% CrI: 0.10–0.16) for cattle and 0.013 (95% CrI: 0.01–0.02) for small ruminants.

**Figure 3:**
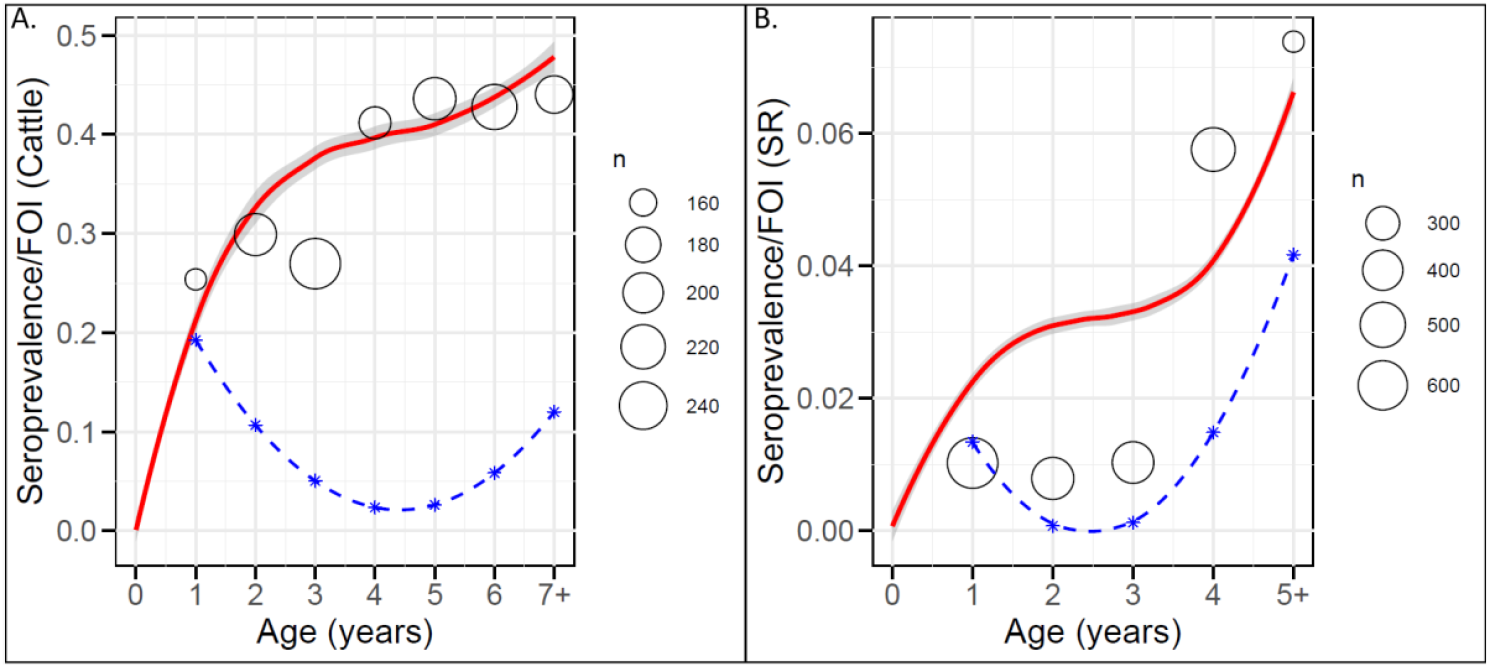
Adjusted RVFV age-stratified seroprevalence in cattle (**A**) and small ruminants (SR) (**B**) in The Gambia, overlaid as circles with size proportional to the number of animals sampled in that age group. The predicted seroprevalence (red line) with 95% CI (grey ribbon) is based on the cumulative FOI from the Grenfell-Anderson model. The age-specific FOI profile based on the Grenfell Anderson’s model (blue dashed line/stars) is shown.

### Risk factors associated with RVFV seropositivity in ruminant livestock Univariable logistic regression analyses

Preliminary univariable analyses identified several potential risk factors associated with RVFV seropositivity in both cattle (Table 2) and small ruminants (Table 3). However, many of these variables did not retain statistical significance in the multivariable glmmLasso models. Household-level management practices, including access to temporary ponds or the floodplains of the Gambia river, appeared to strongly influence RVFV seropositivity in small ruminants.

**Table 2:** Summary statistics and adjusted RVFV seroprevalence for specific predictor variables in the cattle data set. Household-level management practices are based on the six months preceding the study. ^*^ = reference level of categorical variables.

|  |  | Number (%)<br>[Mean, std. dev] | Adjusted Seroprevalence<br>(95% CI) | Odds ratio<br>(95% CI) | P-value |
| --- | --- | --- | --- | --- | --- |
| Individual characteristics |  |  |  |  |  |
| Age (years) | 1 | 156 (10.9) | 25.4 (16.8–33.9) | 1* |  |
|  | 2 | 208 (14.7) | 29.9 (21.2–38.5) | 1.2 (0.7–1.9) | 0.551 |
|  | 3 | 259 (18.3) | 27.0 (19.9–34.0) | 1.3 (0.8–2.1) | 0.284 |
|  | 4 | 170 (12.0) | 41.1 (32.3–50.0) | 1.6 (1.0–2.7) | 0.051 |
|  | 5 | 210 (14.8) | 43.6 (33.4–53.8) | 2.2 (1.6–4.0) | <0.001 |
|  | 6 | 225 (15.9) | 42.8 (33.1–52.4) | 2.0 (1.3–3.2) | 0.003 |
|  | 7+ | 188 (13.3) | 44.0 (32.6–55.4) | 2.3 (1.4–3.7) | 0.001 |
| Sex | female | 968 (68.4) | 39.2 (34.4–43.9) | 1* | <0.001 |
|  | Male | 448 (31.6) | 28.7 (23.3–34.2) | 0.7 (0.5–0.8) |  |
| Household-level characteristics |  |  |  |  |  |
| Have access to the Gambia river floodplains | No | 578 (40.8) | 31.8 (26.8–36.7) | 1* | 0.103 |
|  | Yes | 838 (59.2) | 38.3 (32.4–44.3) | 1.4 (0.9–2.0) |  |
| Transhumant herd movement to access river floodplains | No | 991 (70.0) | 36.1 (31.8–40.3) | 1* | 0.600 |
|  | Yes | 425 (30.0) | 35.0 (23.8–46.2) | 0.9 (0.7–1.3) |  |
| Have access to temporary ponds | No | 112 (7.9) | 33.2 (21.7–44.7) | 1* | 0.845 |
|  | Yes | 1304 (92.1) | 36.1 (31.7–40.6) | 1.1 (0.5–2.1) |  |
| Use vector control strategies | No | 909 (64.2) | 33.9 (28.5–39.3) | 1* | 0.148 |
|  | Yes | 507 (35.8) | 40.0 (34.0–46.0) | 1.2 (0.9–1.7) |  |
| History of abortions | No | 1012 (71.5) | 34.1 (29.0–39.2) | 1* | 0.087 |
|  | Yes | 404 (28.5) | 41.1 (35.0–47.3) | 1.3 (0.9–1.7) |  |
| Cattle co-managed with small ruminants | No | 1243 (87.8) | 36.2 (31.6–40.8) | 1* | 0.244 |
|  | Yes | 173 (12.2) | 33.3 (23.5–43.1) | 0.8 (0.5–1.2) |  |
| Cattle are herded during the dry season | No | 559 (39.5) | 36.7 (31.1–42.3) | 1* | 0.302 |
|  | Yes | 857 (60.5) | 34.1 (28.4–39.8) | 0.9 (0.7–1.1) |  |
| Continuous variables |  |  |  |  |  |
| Total number of cattle in household |  | [44, 27.0] |  | 1.0 (1.0–1.0) | 0.060 |
| Number of new cattle brought into herd |  | [4, 3.1] |  | 1.1 (1.0–1.1) | 0.005 |
| Number of cattle herds in village |  | [7.7, 4.8] |  | 1.0 (0.9–1.0) | 0.431 |
| Distance to water sources (km) |  | [1.6, 0.8] |  | 0.9 (0.7–1.2) | 0.488 |

**Table 3:** Summary statistics and adjusted RVFV seroprevalence for specific predictor variables in the small ruminant (SR) data set. Household-level management practices are based on the six months preceding the study. ^*^ = reference level of categorical variables.

|  |  | Number (%)<br>[Mean, std. dev] | Adjusted Seroprevalence<br>(95% CI) | Odds ratio<br>(95% CI) | P-value |
| --- | --- | --- | --- | --- | --- |
| Individual characteristics |  |  |  |  |  |
| Age (years) | 1 | 633 (29.0) | 0.01 (0–0.03) | 1* |  |
|  | 2 | 434 (19.8) | 0.008 (0–0.02) | 1 (0.5–2.1) | 0.963 |
|  | 3 | 424 (19.4) | 0.01 (0–0.03) | 0.8 (0.4–1.8) | 0.635 |
|  | 4 | 468 (21.4) | 0.06 (0.01–0.1) | 1.8 (0.9– 3.4) | 0.078 |
|  | 5+ | 227 (10.4) | 0.07 (0–0.14) | 1.9 (0.9–4.1) | 0.091 |
| Sex | female | 1691 (68.4) | 3.2 (0.5–5.8) | 1* | 0.614 |
|  | Male | 469 (31.6) | 1.4 (0.6–3.4) | 0.9 (0.5–1.6) |  |
| Household-level characteristics |  |  |  |  |  |
| Have access to the Gambia river floodplains | No | 2094 (95.8) | 2.4 (0.4–4.5) | 1* | 0.001 |
|  | Yes | 92 (4.2) | 17.2 (8.9–43.3) | 5.4 (1.9–15.4) |  |
| Transhumant flock movement to access river floodplains | No | 2101 (96.1) | 2.4 (0.4–4.4) | 1* | <0.001 |
|  | Yes | 85 (3.9) | 16.9 (4.2–37.9) | 5.5 (2.3–12.9) |  |
| Have access to temporary ponds | No | 2028 (88.8) | 2.3 (0.2–4.3) | 1* | <0.001 |
|  | Yes | 158 (11.2) | 13.1 (0.4–25.8) | 5.91 (5.9–5.94) |  |
| Used vector control strategies | No | 1711 (78.3) | 3.1 (0.7–5.5) | 1* | 0.546 |
|  | Yes | 475 (21.7) | 1.3 (0.6–3.3) | 0.8 (0.4–1.5) |  |
| History of abortions | No | 1847 (84.5) | 2.8 (0.6–5.0) | 1* | 0.881 |
|  | Yes | 339 (15.5) | 2.3 (0.05–4.7) | 1.1 (0.5–2.0) |  |
| SR co-managed with cattle | No | 2072 (94.8) | 2.2 (0.2–4.3) | 1* | <0.001 |
|  | Yes | 114 (5.2) | 14.7 (1.0–28.4) | 5.2 (2.5–10.8) |  |
| Continuous variables |  |  |  |  |  |
| Total number of SR in household |  | [14, 7.6] |  | 1.0 (1.0–1.02) | 0.626 |
| Number of new SR brought into herd |  | [7, 3.9] |  | 1.0 (0.9–1.1) | 0.767 |
| Number of SR flocks in village |  | [25.9, 18.4] |  | 1.0 (1.0–1.0) | 0.067 |
| Distance to water source (km) |  | [0.1, 0.5] |  | 1.8 (1.2–2.7) | 0.004 |

### Variable selection in mixed effects multivariable Lasso-penalized GLMMs

The glmmLasso models included eighteen predictor variables collected during the questionnaire survey. We present only the variables selected from the cattle and small ruminant data sets in the main manuscript. The Bayesian Information Criterion (BIC) results as a function of the Lasso penalty parameter (λ), along the glmmLasso model summaries, are provided in the Supplementary Text. In the cattle data set, the optimal penalty parameter (λ_opt_=51.8) shrunk (reduced coefficients to zero) all but eight variables (Figure 4). Due to high correlation of two selected variables, “access to river” was excluded from the final logistic regression model in favor of “transhumance distance to river”. Cattle in the Lower River (OR = 2.2, 95% CI: 1.2–4.0) and North Bank (OR = 2.2, 95% CI: 1.2–4.0) regions had twice the odds of RVFV seropositivity than in the other four regions. Male cattle were 20% less likely to be seropositive (OR = 0.8, 95% CI: 0.6–1.1). Age was included as a continuous predictor, with odds of seropositivity increasing by 13% per year in cattle (OR = 1.13, 95% CI: 1.05–1.21). The total number of cattle in the household had no significant association with seropositivity (OR = 1.0, 95% CI: 1.00–1.02).

**Figure 4:**
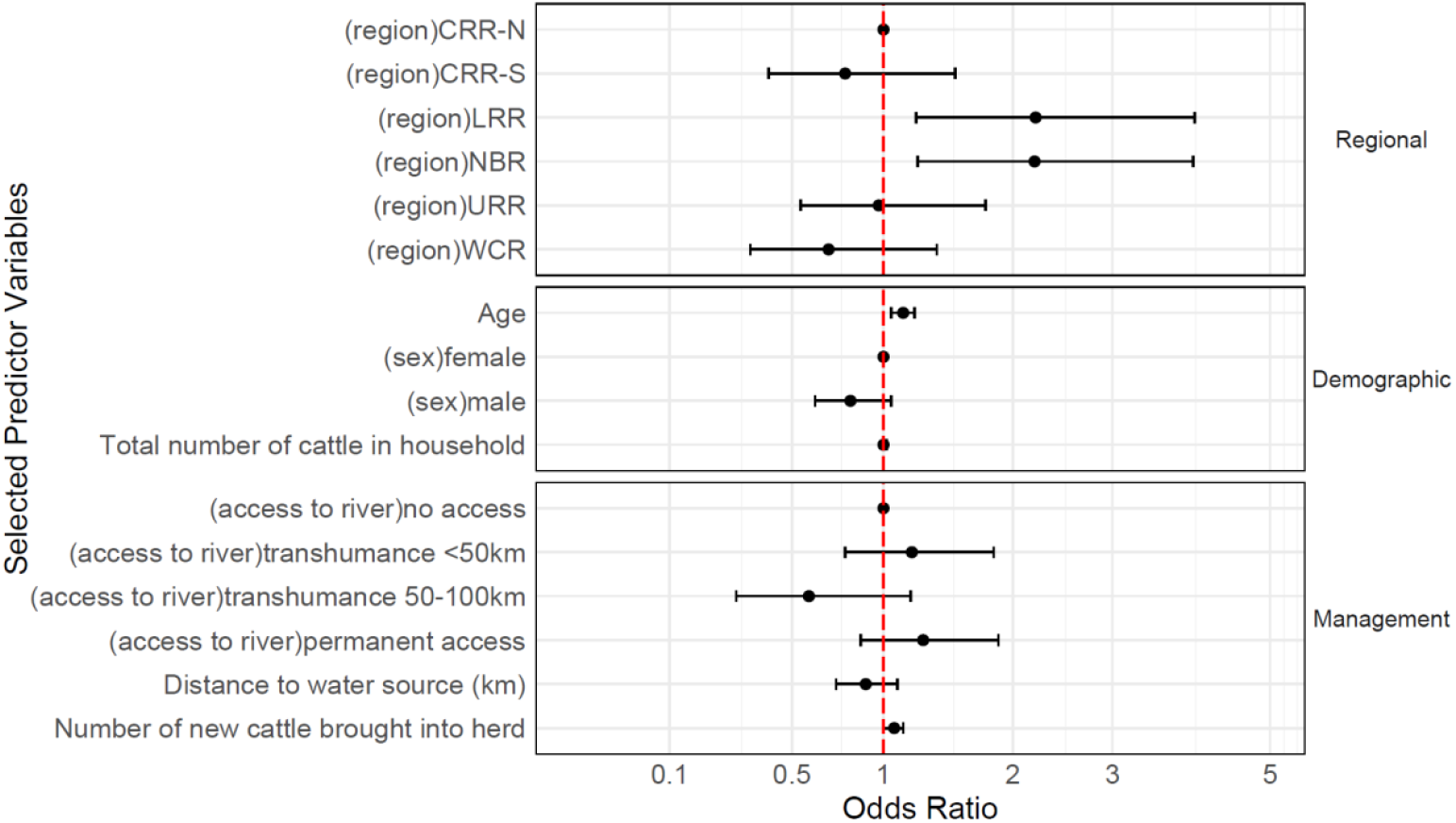
The odds ratios of association between selected predictor variables and RVFV seropositivity in cattle from the glmmLasso analyses. The dots are the estimated odds ratios, and the error bars are the 95% confidence intervals. The random effects of district and village were included in the multivariable logistic regression. The six administrative regions are: CRR-S–Central River Region–South; LRR–Lower River Region; NBR–North Bank Region; URR–Upper River Region; WCR–West Coast Region.

Occupancy of the floodplains of the Gambia river by cattle-owning households (HH) for grazing and water was classified into four categories: (i) year-round occupancy (HH=13) (ii) seasonal access via transhumant movements <50km (HH=46), (iii) seasonal access via transhumant movements covering 50–100km (HH=5), and (iv) no access to the river valley (HH = 68). Cattle that moved within 50km of their homestead to the river valley during the dry season exhibited higher odds of seropositivity (OR = 1.2, 95% CI: 0.8–1.8) compared to those with no access to the river. In contrast, cattle migrating up to 100km to the river valley had lower odds of seropositivity (OR = 0.6, 95% CI: 0.3–1.9), although this result was less definitive due to the small number of long-distance transhumant herds (HH = 5) surveyed. Cattle herds that permanently occupy the floodplains of the Gambia river exhibited approximately 30% higher odds of seropositivity (OR = 1.3, 95% CI: 0.9–1.9) compared to those with no access.

Introduction of new cattle into a household herd was associated with 7% greater odds of seropositivity (OR = 1.07, 95% CI: 1.0–1.1). Seropositivity was negatively associated with proximity to the water sources used by the cattle herd. Specifically, for every kilometer increase in distance to water, the odds of seropositivity in cattle decreased by 10% (distance to water source: OR = 0.9, 95% CI: 0.7–1.1).

The glmmLasso procedure (λ_opt_=30.3) applied to the small ruminant data set selected three variables (Figure 5). Access to temporary ponds during the wet season was associated with a threefold increase in the odds of seropositivity in small ruminants (OR = 3.2, 95% CI: 1.0– 10.5). Small ruminants that are kept and co-managed alongside cattle herds (OR = 1.8, 95% CI: 0.5–6.3), and those involved in seasonal transhumant movement (OR = 1.4, 95% CI: 0.4– 4.8) had higher odds but non statistically significant association with seropositivity.

**Figure 5:**
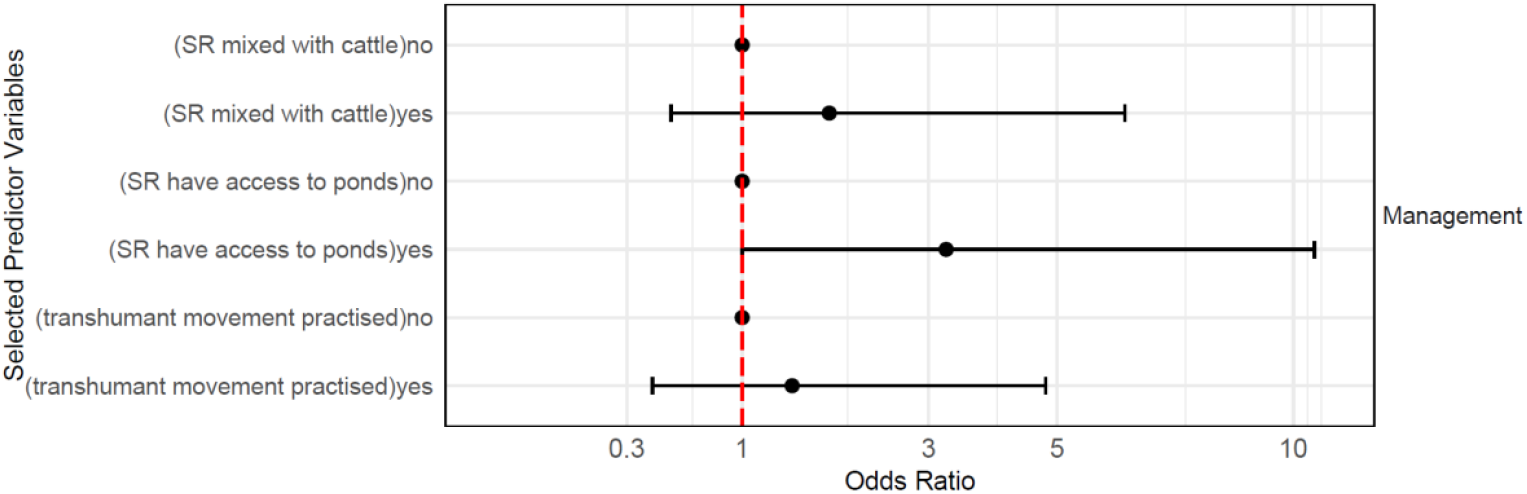
The odds ratios of association between selected predictor variables and RVFV seropositivity in small ruminants from the glmmLasso analyses. The dots are the estimated odds ratios, and the error bars are the 95% confidence intervals. The random effects of district and village were included in the multivariable logistic regression. The six administrative regions are: CRR-S–Central River Region–South; LRR–Lower River Region; NBR–North Bank Region; URR– Upper River Region; WCR–West Coast Region.

## Discussion

This study provides insights into the sero-epidemiology of RVFV in The Gambia, offering the first population-level RVFV seroprevalence estimates and identifying risk factors associated with seropositivity among ruminant livestock. To the best of our knowledge, no confirmed RVF cases have been documented in ruminants since the last reported outbreak in 2002. We observed substantial variation in RVFV seropositivity across study villages, suggesting spatial heterogeneity in exposure intensity, with the Gambia river acting as a critical hotspot for livestock infection. Consistent with studies in Kenya [66], Mauritania [67], and South Africa [33], RVFV seroprevalence in The Gambia was higher in cattle (36.8%) compared to goats (2.1%) and sheep (4.3%). However, the difference in seroprevalence between cattle and small ruminants (sheep and goats) is more marked in The Gambia than reported in most other studies. While factors such as species-specific susceptibility [68], vector feeding preferences [69] and body size may be plausible explanations for this observation, specific features of livestock grazing and management in The Gambia which affect differential exposure to RVFV-competent mosquitoes may be notable contributing factors. For example, a greater proportion of cattle herds (59.2%) than small ruminant flocks (4.2%) in our study permanently occupied or seasonally accessed the floodplains of the Gambia river, and thus spent more time in areas that are eco-climatically favorable for mosquito vectors.

The glmmLasso models further highlighted the role of the river in the epidemiology of RVFV in The Gambia, with transhumant livestock movements to its floodplains associated with higher odds of exposure for both cattle and small ruminants. Thirty percent (30%) of the surveyed cattle herds moved into the river floodplains during the dry season, and all the small ruminants practicing transhumance (3.9%) were co-managed with the cattle herds during seasonal movements. These movements concentrate livestock in riverine ecosystems where entomological studies have caught thriving populations of mosquitoes, such as *Culex poicilipes* and *Mansonia africana*, naturally infected with RVFV [70, 71]. Previous studies have observed that regions in the Sahelian zone, particularly those near large rivers and lakes, are highly suitable for RVF epidemics and have high observed seroprevalence during the IEP [24, 72, 73]. Fifty-four mosquito species and varieties have been described for The Gambia [74], including RVFV-competent *Cx. poicilipes, Cx. tritaeniorhynchus, Ma. africana*, and *Ma. uniformis*. A dry-season study in irrigated rice fields adjacent the Gambia river found *Ma. africana* and *Ma. uniformis* present throughout a 21-week study, while *Culex* spp. peaked four weeks after irrigation began [75]. The presence of RVFV-competent mosquitoes along the Gambia river, combined with a substantial ruminant population in the dry season suggests an environment conducive for year-round virus circulation and is likely a significant risk factor for RVFV seropositivity in the region.

It is noteworthy that the seroprevalence recorded for cattle in The Gambia was substantially higher than has been reported in many other regions where RVF is considered endemic or hyperendemic, and in studies that have adopted the same IDScreen_®_ RVF Multi-species c-ELISA diagnostic assay (e.g. 7.3% in Egypt [76] and 4.4% in Tanzania [77]). Although RVF has not typically been considered a major human or livestock health problem in The Gambia, these results clearly indicate the potential for zoonotic transmission of RVFV, given the close interaction between humans and livestock in the country. Additionally, the impacts on cattle health and productivity cannot be ruled out. Despite the high seroprevalence, there have been no reports of large-scale abortion events or neonatal mortalities among livestock in The Gambia since 2002. This may be attributed to limited surveillance infrastructure or under-reporting of cases to authorities – common challenges across sub-Saharan Africa [78]. While our analysis found no association between RVFV seropositivity and abortion history, other etiologies of animal abortions [79] should be considered. The N’Dama breed of cattle sampled in this study is well known for its tolerance to diseases such as trypanosomosis [80], and may exhibit a similar resilience to RVFV infection. A further possibility is that transmission routes in The Gambia expose cattle to low infective doses that result in only subclinical signs. Lastly, it is also possible that the RVFV strain(s) circulating in The Gambia may be less virulent than those responsible for outbreaks in neighbouring Mauritania and Senegal.

While high seropositivity could result from epidemic transmission, our analysis of age-seroprevalence data supported an interpretation of endemic or hyperendemic circulation of RVFV in The Gambia. Seropositivity increased with age and seropositive individuals were detected in the youngest age groups of both cattle and small ruminants. The predicted seroprevalence curve based on the FOI estimates of the quadratic model for cattle showed a rapid increase in infections during calfhood, followed by a plateau by early adulthood, a pattern consistent with other diseases [81]. Our age-independent FOI estimate of 0.12 in cattle and 0.013 in small ruminants in this study exceed estimates among cattle surveyed in Cameroon [25] and Madagascar [82], and small ruminants in Tanzania [77], indicating relatively high transmission rates in The Gambia. Although these FOI estimates are based on a suboptimal model, they provide an important baseline for future epidemiological studies of RVFV in The Gambia.

The abundance of RVFV-competent mosquitoes around temporary ponds is consistently reported to facilitate RVFV transmission in West Africa [23, 73, 83]. In our study, small ruminants accessing these ponds had significantly higher odds of seropositivity, supporting findings from Senegal that linked the combination of temporary ponds and vegetative cover to increased seropositivity in small ruminants [84]. Eight RVFV-competent *Aedes* mosquito species, including *Ae. vexans* and *Ae. ochraceus*, have been identified from breeding sites near temporary rivers in Senegal [21], but their roles in transovarial transmission remain unconfirmed, although not completely ruled out. *Aedes* spp. may initiate RVFV transmission cycles around rain-fed temporary ponds that are then amplified in the latter phase of the wet season by *Culex, Anopheles*, and *Mansonia* mosquito species. While RVFV-competent mosquitoes from these genera have been reported in The Gambia [74, 75], further studies are needed to clarify their roles in local RVFV dynamics.

The introduction of new animals was selected as an important risk factor for RVFV seropositivity, with 29% of households reporting the introduction of new ruminants in the six months preceding the study. The Gambia maintains an unrestricted transboundary trade of ruminants, including from RVFV-endemic regions of Mauritania and Senegal to meet a high demand for these ruminants during the Muslim festive periods of Eid. This risk of RVFV introduction to the local livestock population is similar to explanations given for the outbreak in Saudi Arabia in 2000 and Mayotte in 2007, where phylogenetic analyses linked virus entry to importation of asymptomatic but infectious ruminant from the Horn of Africa [85, 86]. To mitigate this risk, additional educational campaigns that promote biosecurity measures, such as quarantine protocols for new livestock, should be prioritized. Effective strategies could include community-based training, radio messaging, farmer-to-farmer knowledge sharing, and strengthening engagement with the veterinary service.

Proximity to water was a risk factor for RVFV seropositivity in cattle, with each additional kilometer distance to water sources reducing the odds of seropositivity by 10%. Cattle tethered outdoors overnight near water sources are particularly vulnerable to nocturnal *Culex* spp. and crepuscular *Aedes* spp. mosquitoes [87, 88]. Proximity to intermittent flooded areas or permanent freshwater swamps has been identified as a strong predictor of RVF outbreaks in sub-Saharan Africa [24]. Previous RVF sero-surveillance studies also associated higher RVFV seropositivity to proximity to major water bodies, such as the Senegal river [89] and Lake Malawi [90]. In contrast, no significant association was observed between proximity to water and RVFV seropositivity in small ruminants, potentially due to the small number of seropositives in our study. However, this finding aligns with research from Senegal, where distance to temporary ponds did not have a protective effect on RVFV exposure in small ruminants [23].

The study also revealed differences in seroprevalence among male and female cattle. The cause of this sex-based variation in RVFV exposure in The Gambia is unclear, as male and female ruminants are generally managed together. However, the 2016 Livestock Census indicates that females constitute approximately 70% of the cattle population in The Gambia. As a high-risk area, a RVF outbreak affecting the larger female population could have severe consequences on the livelihoods of vulnerable families, many of whom rely on milk for sustenance and depend on the reproduction of their livestock for wealth accumulation [91].

We used the penalized regression approach to more parsimoniously select important predictor variables (compared to traditional approaches, e.g., stepwise selection), through iterative estimation of the penalty parameter [60]. Our approach mitigated the risk of overfitting, and provided BIC measures to assess model fit [92]. Here, the glmmLasso algorithm demonstrated high accuracy in variable selection and computational efficiency.

## Conclusion

This study presents the first comprehensive analysis of RVFV seroprevalence among ruminant livestock in The Gambia, revealing a significantly higher seroprevalence in cattle compared to small ruminants. Our findings suggest endemic circulation of RVFV, with seropositivity increasing with age and observed in the youngest age groups. The low seropositivity in small ruminants indicates limited herd immunity and therefore potential for large outbreaks when conditions become favourable for virus amplification. We highlight the role of the floodplains of the Gambia river in increasing the risk of RVFV seropositivity, alongside specific herd management practices including transhumant movements, introduction of new animals, proximity to water sources, and access to temporary ponds as key risk factors. These findings underscore the need for preventive measures, such as topical insecticide treatment or vaccination, to reduce infection risk prior to transhumant movements into the floodplains. Despite the absence of clinical outbreaks, the high seroprevalence stresses the importance of establishing active surveillance systems to monitor livestock abortions and unexplained neonatal mortalities, and to enable early detection and effective management of potential RVF outbreaks.

## Acknowledgements

We thank the communities and livestock-owning households for participating in the study. We are also deeply grateful to the livestock assistants of the Department of Livestock Services in The Gambia for their contributions to the field work.

## Author Contributions

**E.J**., **D.T.H**. and **S.C**. conceptualized the study; **E.J**. acquired the funding; **E.J**., **Y.B**., **O.S**., **A.S**. and **K.D**. coordinated and administered the field work; **E.J** and **A.M**. conducted the laboratory investigations; **E.J**. conducted the data analyses & visualization; **E.J**. wrote the original draft with input from **D.T.H**. and **S.C.; D.T.H**. and **S.C**. supervised the project; All authors reviewed, edited and approved the manuscript.

## Data Availability Statement

The data that support the findings of this study are available on request from the corresponding authors. The data are not publicly available due to privacy or ethical restrictions.

