## Supplementary Information for "Sero-epidemiology of Rift Valley fever virus in ruminant livestock in The Gambia"

### Supplementary Text

### Intracluster correlation coefficients

One of the main strengths of our study is the use of simulation-based power analysis, which accounted for the impact of random effects into our study design. This approach effectively captured the variability in seroprevalence across the 52 study villages was, which appeared to be largely influenced by livestock management practices. Notably, the ICCs of both species groups (0.13 for cattle and 0.22 for small ruminants) highlight that herd level factors contribute moderately to the variability in seroprevalence. This finding underscores the need for surveillance programmes to account for clustering effects to accurately assess and interpret RVF dynamics, and ensure surveillance efforts and resource allocation can be more targeted to herds with higher risk of exposure (those with specific livestock management practices). These ICC values align with those reported in endemic settings in Kenya (0.30) (Bett *et al.*, 2018) and South Africa (0.22 – 0.41) (Bett *et al.*, 2018; Ngoshe *et al.*, 2020), suggesting shared epidemiological patterns across African regions with different ecological and livestock management practices.

### Variable selection in glmmLasso

Briefly, we started by computing the control starting values, Delta and Q, using glmmPQL (Venables and Ripley, 2002). We hand-coded lists to save the Delta_start vector of the likelihood-functions for the fixed and random effects, and the Q_start matrix for the random effect variance-covariances computed for specific values of the LASSO penalty term (λ) in the first model fit. These values were recovered in the final re-estimation fit.

To estimate the optimal λ value, we first fitted a starting model scanning through a sequence of values ($0 \leq \lambda \leq 500$) in 100 steps such that all predictor coefficients are reduced to zero (shrunk to zero). The optimal λ was chosen by identifying the model with the lowest Bayesian Information Criterion (BIC). As recommended by Groll and Tutz (2014), a final glmmLasso model was re-fitted by simple Fisher scoring using estimates of the starting values obtained from the previous fit.

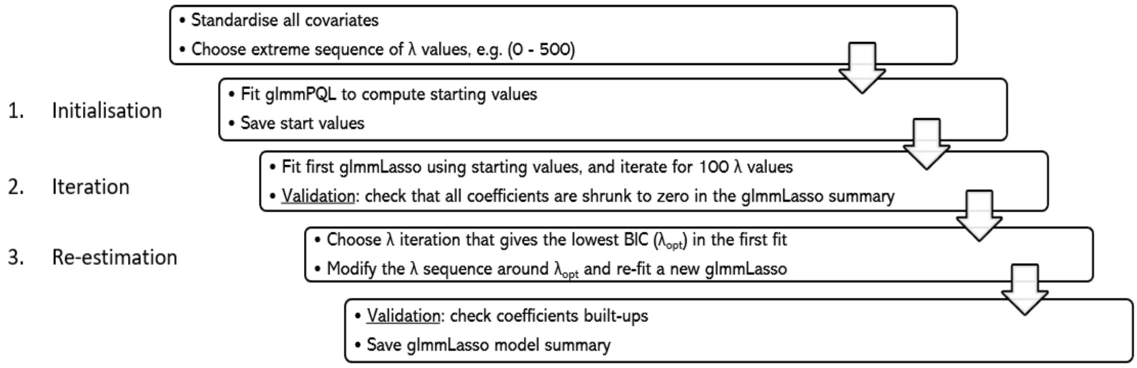


*Figure S1: Flowchart of the glmmLasso procedure used in the analyses. The boxes represent the main steps along the L1-regularisation path to selecting the predictor variables most important in explaining RVFV seropositivity in ruminants in The Gambia.*

### glmmLasso models

### Cattle data set

The glmmLasso model included 18 predictor variables collected during the questionnaire survey. In the initialisation step, the penalty sequence (0 – 500) conferred a very large penalisation (and degree of shrinkage) such that all grouped and standardised covariate coefficients were reduced to zero. To correct the glmmLasso predictive power, several controlled changes were computed in the values of λ until λ = 1 – 180. The glmmLasso model output converged and the coefficient estimates did not change with λ values between 180 – 200. The optimal λ ($\lambda_{opt}$) of the glmmLasso procedure for the cattle data set was 51.82, corresponding to the model with the lowest BIC (Figure S2).

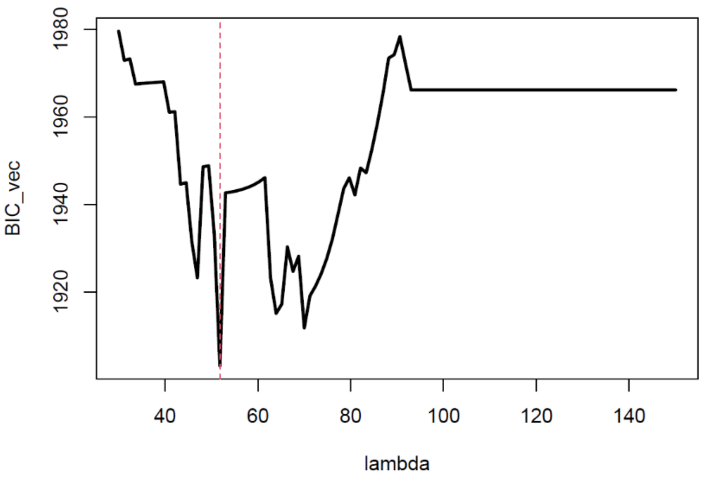


*Figure S2: BIC results for the glmmLasso as a function of the λ sequence for the cattle data set; imposed red dotted vertical line shows the optimal λ value.*

The $\lambda_{opt}$ shrunk (reduced exactly to zero) all but seven variables, which suggested a strong association between these maintained non-zero variables and RVFV seropositivity (McNeish, 2015). The final glmmLasso model output is presented in Table S1.

*Table S1: Model summary from the glmmLasso approach of the cattle data set showing coefficient estimates of the selected variables. HH = household. The six administrative regions are: crr-s – Central River Region–South; lrr – Lower River Region; nbr – North Bank Region; urr – Upper River Region; wcr – West Coast Region.*

| Label | Estimate | StdErr | z.value | p.value |
|---|---|---|---|---|
| (Intercept) | -0.7449 | 0.0965 | -7.7171 | 0.0000 |
| as.factor(region)crr-s | -0.2751 | 0.2755 | -0.9986 | 0.3180 |
| as.factor(region)lrr | 0.7774 | 0.2727 | 2.8507 | 0.0044 |
| as.factor(region)nbr | 0.7773 | 0.2940 | 2.6434 | 0.0082 |
| as.factor(region)urr | -0.0235 | 0.3868 | -0.0608 | 0.9516 |
| as.factor(region)wcr | -0.3666 | 0.5190 | -0.7064 | 0.4800 |
| age | 0.2399 | 0.0647 | 3.7056 | 0.0002 |
| as.factor(sex)male | -0.2070 | 0.1403 | -1.4751 | 0.1402 |
| as.factor(catttle.mixed.with.other.ruminants)yes | 0.0000 | | | |
| average.rainfall.in.district | 0.0000 | | | |
| as.factor(brought.new.cattle.into.HH)yes | 0.0000 | | | |
| as.factor(vector.control.practised)yes | 0.0000 | | | |
| total.number.of.cattle.in.HH | 0.0255 | 0.0758 | 0.3364 | 0.7365 |
| as.factor(any.cattle.died.of.disease)yes | 0.0000 | | | |
| as.factor(cattle.have.access.to.river)yes | 0.3401 | 0.4030 | 0.8439 | 0.3987 |
| as.factor(cattle.have.access.to.ponds)yes | 0.0000 | | | |
| as.factor(contact.with.30.other.herds)yes | 0.0000 | | | |
| as.factor(transhumant.movement.practised)yes | 0.0000 | | | |
| as.factor(transhumance.distance.to.reach.river)short.distance | -0.1281 | 0.2176 | -0.5887 | 0.5561 |
| as.factor(transhumance.distance.to.reach.river)long.distance | -0.8301 | 0.3886 | -2.1360 | 0.0327 |
| as.factor(transhumance.distance.to.reach.river)river-residents | -0.0839 | 0.5339 | -0.1572 | 0.8751 |
| distance.to.water.sources.km | -0.0708 | 0.0792 | -0.8939 | 0.3714 |
| as.factor(any.cattle.abortion.stillborn)yes | 0.0000 | | | |
| number.of.cattle.brought.into.herd | 0.1530 | 0.0723 | 2.1174 | 0.0342 |
| number.of.cattle.herds.in.village | 0.0000 | | | |

### Small ruminant data set

As above, several controlled changes were computed in the values of λ until λ = 1 – 100. The glmmLasso model output converged and the coefficient estimates did not change with λ values between 70 – 150. The optimal λ ($\lambda_{opt}$) of the glmmLasso procedure for the small ruminant was 30.30, corresponding to the model with the lowest BIC (Figure S3).

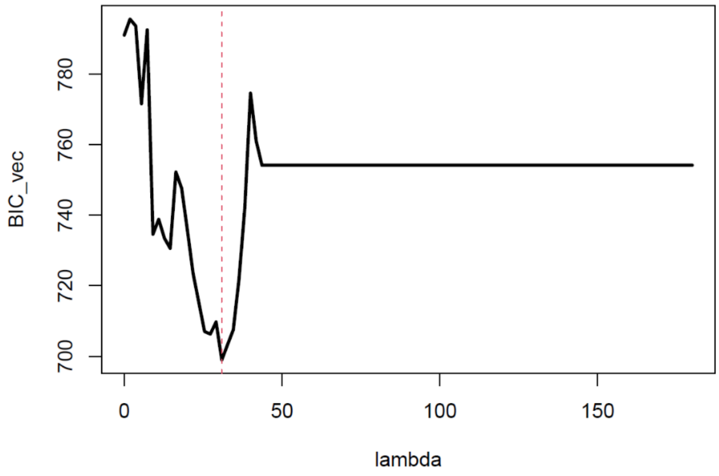


*Figure S3: BIC results for the glmmLasso as a function of the λ sequence for the small ruminant (SR) data set; imposed red dotted vertical line shows the optimal λ value.*

The glmmLasso procedure selected only three variables in the SR data set, which suggested a strong association between these maintained non-zero variables and RVFV seropositivity (McNeish, 2015).

*Table S2: Model summary from the glmmLasso approach of the small ruminant (SR) data set showing coefficient estimates of the selected variables. HH =*

*household. The six administrative regions are: crr-s – Central River Region–South; lrr – Lower River Region; nbr – North Bank Region; urr – Upper River Region; wcr – West Coast Region.*

| Label | Estimate | StdErr | z.value | p.value |
|---|---|---|---|---|
| (Intercept) | -3.7867 | 0.2552 | -14.8371 | 0.0000 |
| as.factor(region)crr-s | 0.0000 | | | |
| as.factor(region)lrr | 0.0000 | | | |
| as.factor(region)nbr | 0.0000 | | | |
| as.factor(region)urr | 0.0000 | | | |
| as.factor(region)wcr | 0.0000 | | | |
| age | 0.0000 | | | |
| as.factor(sex)male | 0.0000 | | | |
| as.factor(SR.mixed.with.other.ruminants)yes | 0.4317 | 0.6407 | 0.6739 | 0.5004 |
| as.factor(brought.new.SR.into.HH)yes | 0.0000 | | | |
| as.factor(vector.control.practised)yes | 0.0000 | | | |
| total.number.of.SR.in.HH | 0.0000 | | | |
| as.factor(any.SR.died.of.disease)yes | 0.0000 | | | |
| as.factor(SR.have.access.to.river)yes | 0.0000 | | | |
| as.factor(SR.have.access.to.ponds)yes | 1.2822 | 0.6270 | 2.0450 | 0.0409 |
| as.factor(contact.with.30.other.flocks)yes | 0.0000 | | | |
| as.factor(transhumant.movement.practised)yes | 0.3564 | 0.6209 | 0.5741 | 0.5659 |
| as.factor(transhumance.distance.to.reach.river)short.distance | 0.0000 | | | |
| as.factor(transhumance.distance.to.reach.river)long.distance | 0.0000 | | | |
| distance.to.water.sources.km | 0.0000 | | | |
| as.factor(any.SR.abortion.stillborn)yes | 0.0000 | | | |
| number.of.SR.brought.into.flock | 0.0000 | | | |

### Limitations

Some limitations should be noted when interpreting these RVFV seroprevalence and FOI estimates. The detection of anti-RVFV IgG antibodies in serum samples depended on the use of a commercially available assay. This assay has been validated in a veterinary reference laboratory in neighbouring Senegal, and another performance study in Cameroon found it to be highly specific for RVFV (de Bronsvoort *et al.*, 2019). The questionnaire data collected during the cross-sectional survey are limited by recall and social desirability bias and may misrepresent the associations between these risk factors with RVFV seropositivity. While dentition is a useful way of aging ruminants, variations in the nutrition, environment and well-being of ruminants possibly make consistent estimation of age difficult. This variability in age classification may then constrain our estimation of age-dependent rates such as the FOI.
